# Emergent Leadership Driven by Asymmetric Stability Shapes Learning in Dyadic Motor Coordination

**DOI:** 10.1101/2025.05.30.657019

**Authors:** Xiaoye Michael Wang, Geoffrey P. Bingham, Qin Zhu

**Author notes:** Corresponding author: Xiaoye Michael Wang.

## Abstract

Coordinated actions are fundamental to human behavior. Even without explicit communication, stable coordination patterns and implicit leadership roles can emerge. This study examines the dynamic nature of leadership and its impact on perceptuomotor learning in a dyadic coordination task. Paired participants (dyads) controlled targets using joysticks to produce a novel coordination pattern, which was learned using visual feedback without direct communication over multiple sessions. Results showed that leadership was an emergent property of the system, dynamically shaped by the interplay between individual and system-level stability. At the system level, leadership role transitions modulated the dyad’s learning, characterized by distinct learning models. At the individual level, leaders maintained overall stability but exhibited occasional bursts of instability to assert control and reinforce coordination. These findings highlight the role of leader-follower dynamics in learning and performance, showing that leadership is both a limiting factor and a facilitator for coordination.

## Introduction

The ability of individuals, be they cells, animals, or humans, to coordinate their actions underlies many complex adaptive behaviors, from tissue development to flocks of migrating birds and collaborative human actions^1–3^. A central question is how such coordination is achieved, especially during direct interactions to achieve a common goal^4^. Dyadic coordination between two people, the simplest form of collective action, is critical for investigating this question^5^. Often, these interactions are not symmetrical: Individuals frequently adopt specialized leader or follower roles to improve the efficiency and robustness of the joint outcome^6–8^. Therefore, identifying the underlying mechanisms that govern the formation and function of these dynamic roles is essential for a deeper understanding of social coordination. Previous research suggests internal states, such as energetic reserves, can dictate which individual in a foraging pair initiates movement^6^. Individual behavioral traits or personalities, like boldness, have also been shown to correlate with leadership tendencies^9^. Beyond individual attributes, social dynamics play a critical role. Dominance hierarchies^7^, the social construction and negotiation of identities^8^, and even implicit cognitive mechanisms^10^ could influence who leads and who follows. Once roles are established, specific interaction strategies (e.g., kinematic signaling^11^) or dynamic switching between compromise and leadership based on navigational conflict^12^ help maintain coordination.

While earlier studies identify factors correlating with leadership or specific coordination tactics, they often rely on static individual characteristics (states, traits), pre-defined social structures, or cognitive frameworks. Less understood are the dynamic behavioral signatures that might intrinsically define leader and follower roles as they unfold during an interaction. Can the temporal structure of behavior itself, reflecting movement variability, provide a signature of functional roles independent of internal states or explicit signaling? Crucially, a significant knowledge gap also exists in linking how such dynamic roles are enacted to their functional consequences for the dyad. Specifically, how do different emergent leadership dynamics influence the partnership’s ability to adapt or learn over time?

In this study, we investigated how leader-follower relationships emerge in dyadic rhythmic coordination and how they influence perceptuomotor learning of a novel coordination pattern. Coordinated rhythmic movements require limb segments or targets to oscillate while maintaining a specific relative phase^13–18^. While in-phase (0°) and antiphase (180°) coordination occur naturally, achieving a 90° relative phase requires explicit perceptuomotor training^19–26^. In a dyadic context, each participant independently controls a target to achieve the desired coordination. When errors occur, both individuals must correct their movements, but simultaneous adjustments and/or over-correction would interfere with coordination. Since 90° coordination demands a fixed relative phase at a constant movement frequency^27,28^, the optimal strategy involves a leader setting the tempo while a follower adapts limb kinematics. Thus, training could help dyads establish and sustain such leader-follower dynamics. Without explicit instructions or direct communication, participants formed dyads and trained over multiple sessions. We examined how leadership roles emerged over time and how shifts in these roles influenced coordination stability and learning outcomes. We hypothesized that leadership would not be a fixed trait, but an emergent property shaped by interaction dynamics. Specifically, we expected stable leadership roles to facilitate learning, whereas ineffective coordination would trigger role reversals. By analyzing these processes, we aimed to uncover fundamental principles of spontaneous leadership in dyad coordination and its role in perceptuomotor learning.

## Results

Fifteen pairs of participants performed a dyadic unimanual coordination task in which each individual used a joystick to independently control the lateral position of one of two white dots on a computer screen (Figure 1a). Each participant controlled a moving dot, oscillating it back and forth across the screen, while coordinating with their partner to maintain a 90° relative phase between the dots’ movements (Figure 1b). Each dyad went through a series of training sessions across multiple days, in which individuals received continuous visual feedback of the movement dots on the screen to facilitate learning while not being allowed direct visual or verbal communication with one another. A metronome set to 0.75 Hz provided auditory guidance for the first half of each trial, whereas the dots on the screen changed color to indicate whether the dyad’s relative phase was within the target range (90° ± 20°). Kinematic data from the joystick movements were recorded at 120 Hz and analyzed using TAT-HUM^29^, a Python-based kinematic analysis toolkit. The phase of each dot was determined as the angle of the velocity-position vector in the phase plane, unwrapped around 360° to ensure continuity. Relative phase is the phase difference between the two dots folded at 180°, resulting in values between 0° and 180°. Performance was measured by the proportion of time on task (PTT), determined as the proportion of time in which the relative phase was ±20° within the desired relative phase of 90°. Training was terminated if the PTT exceeded 60% in a session.

**Figure 1.**
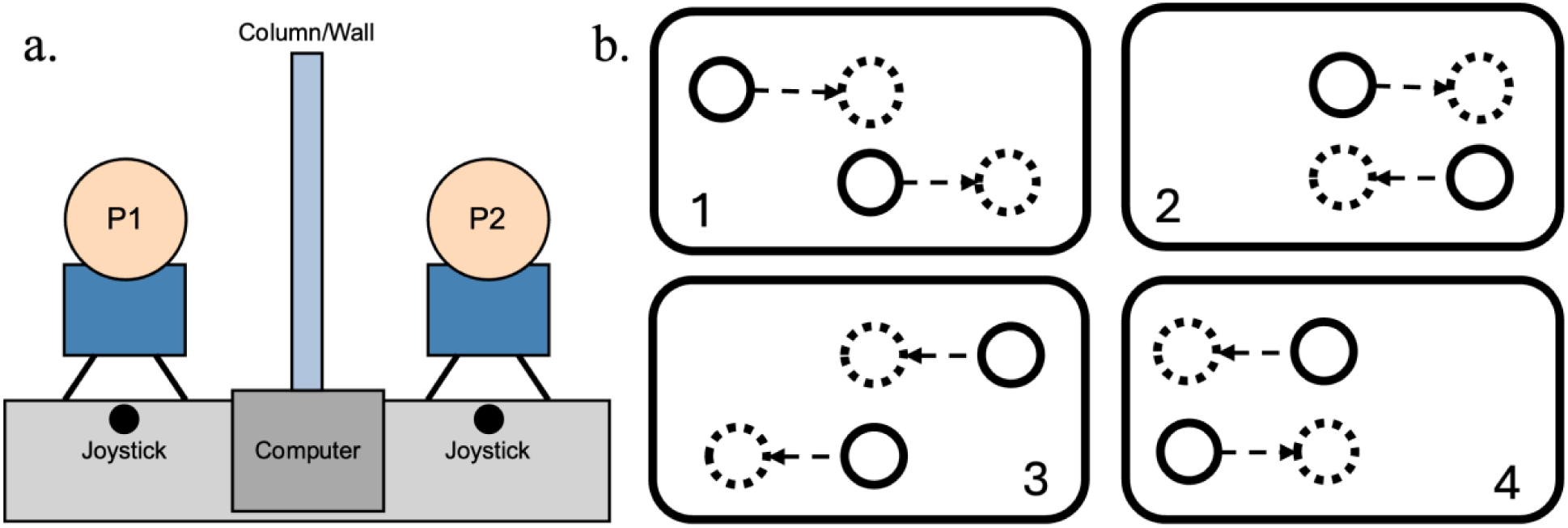
(a) Experimental setup of the training and testing sessions for each dyad. Individuals from a dyad sat next to each other separated by a wall so that they could not see each other, although both could see the computer screen. Each individual independently controls a joystick that moves a dot displayed on the computer. (b) A chronological depiction of the pattern of 90° coordination. The solid circles indicate the targets’ current position whereas the dotted circles indicate the targets’ position at the following segment to produce 90° relative phase.

### Leadership in Training

To identify the leader-follower relationship, the phase difference between the top and bottom dots was analyzed. The unwrapped phase of each dot represents its position in the movement cycle (Figure 2a), and the sign of the phase difference indicates which dot is ahead or behind (Figure 2b). In this context, the leader is the partner whose movements are consistently ahead in the movement cycle, while the follower is the one whose movements lag behind, continuously adjusting to “catch up” with the leader. A series of one-sample t-tests were used to compare the phase difference with 0. A significantly positive phase difference indicates that the top dot was the leader, whereas a negative phase difference indicates that the bottom dot was the leader. Across all training sessions and dyads, there were only 18 trials (or 0.25%) in which neither individual in the dyad emerged as the leader, which were removed from subsequent analysis.

**Figure 2.**
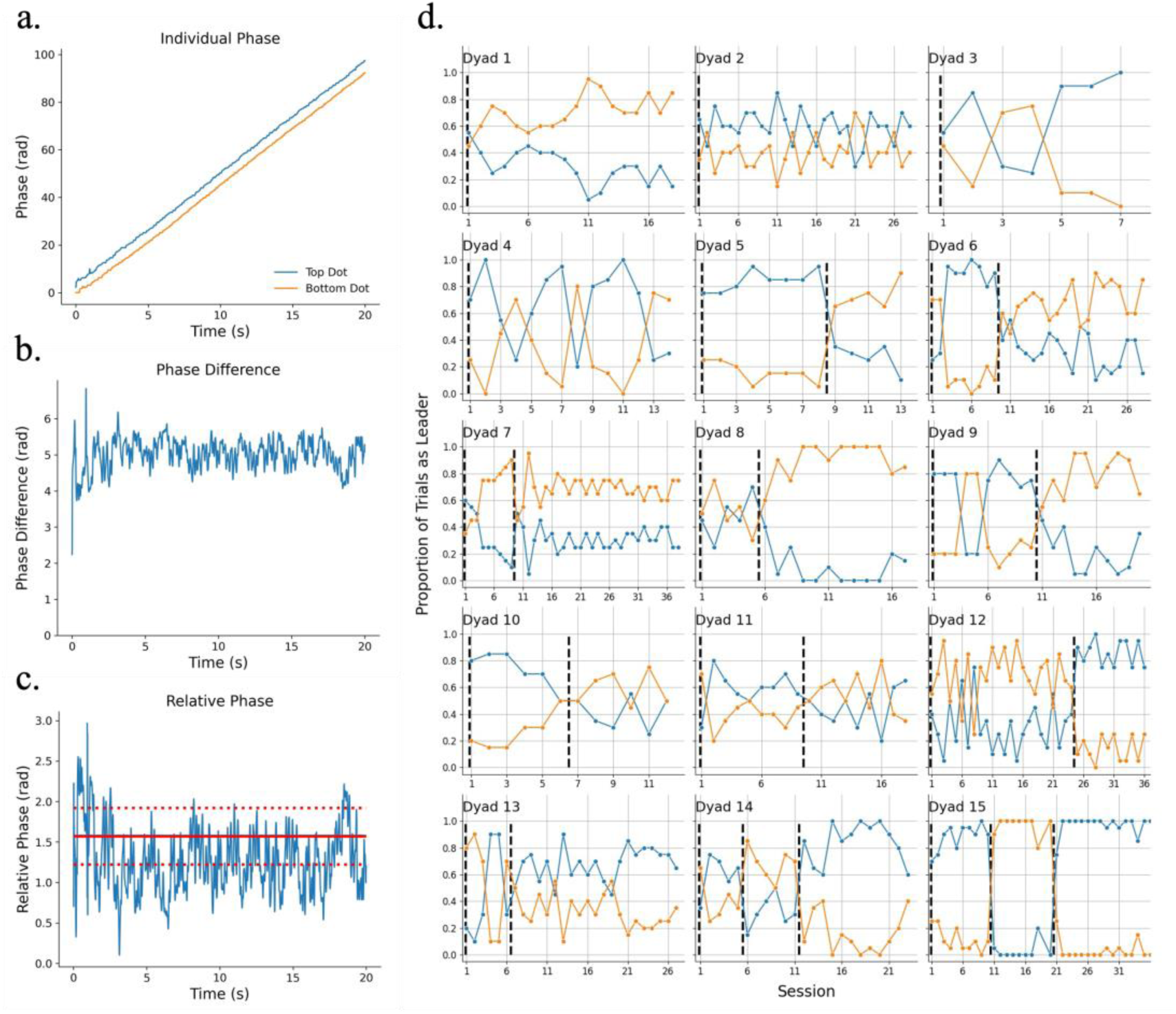
(a) The phases for individuals in a dyad in a single trial as a function of time. The phase for each individual was unwrapped around 2*π* (360°) to yield a continuous phase representation. In this example, the magnitude of the top dot’s phase is consistently higher than that of the bottom dot’s, indicating the top dot had been consistently leading. (b) The phase difference between the two individuals. The positive phase difference again indicates the top dot had been leading the bottom dot in this trial. (c) The relative phase between the two individuals, which is their phase difference folded at *π* (180°). The red solid line and the dotted lines indicate the target relative phase *π*/2 (or 90°) and the error bands *π*/2 ± *π*/9 (or 90° ± 20°), respectively. The proportion of time on task (PTT) is the proportion of samples that fall within the error bands for each trial. (d) The proportion of trials in which the top (blue) and bottom (orange) dots were leading, plotted across training sessions for each dyad. The black dashed lines indicate the beginning of a training segment where one individual has consistently led across multiple sessions. To account for short-term fluctuations, a switch in leadership was only considered a new segment if it lasted for three or more consecutive sessions.

Different dyads exhibited distinct leadership dynamics throughout training (Figure 2d). For some dyads, one individual consistently led all sessions (e.g., Dyad 1), despite occasional brief switches in the leader-follower relationship that lasted only one or two sessions before reverting to the original roles (e.g., Dyad 4). In contrast, some dyads showed a clear leadership transition during training with different individuals leading for extended periods across multiple segments (e.g., Dyad 5). To examine how leadership dynamics influenced dyadic coordination and motor learning, training sessions were segmented based on periods where a single individual remained as the leader. To account for short-term fluctuations, a switch in leadership was only considered a new segment if it lasted for three or more consecutive sessions. For example, for Dyad 3, the bottom dot led for two consecutive sessions in the middle of training, but since this was fewer than three sessions, the entire training period was treated as a single segment. This segmentation method revealed three distinct leadership patterns:

- Scenario 1, **Persistent Leadership** (Dyads 1-4): One individual led throughout training, despite occasional short-term fluctuations. In Dyads 3 and 4, brief role changes occurred but never lasted more than three consecutive sessions.
- Scenario 2, **Single Role Switch** (Dyads 5-13): One individual led during the first half of training, after which leadership switched to the other person for the remainder of the sessions.
- Scenario 3, **Double Role Switch** (Dyads 14-15): Leadership switched twice, where one individual initially led, then the other took over, before leadership reverted to the original leader.

### Leadership Dynamics and Dyad Learning

Different learning models, including linear, power law, logistic growth, and exponential decay, were fitted to the PTT of each training segment to evaluate the effect of leadership dynamics on dyad learning. The best-fitting model was selected based on the lowest Bayesian information criterion (BIC) score (Figure 3a, Table 1). Overall, training segments based on the leader-follower relationship captured variations in learning patterns across sessions. In Scenario 1 (Persistent Leadership; Dyads 1-4), a single segment adequately represented leadership dynamics throughout training. While the best-fitting learning model varied across dyads, all fitted models were statistically significant with relatively high *R*^2^, indicating consistent improvement in performance. Notably, most dyads exhibited rapid initial improvement that later plateaued, as reflected in power law and exponential models (except for Dyad 4, which followed a linear trajectory). This suggests that dyads in this scenario experienced early gains in coordination efficiency that could be attributed to the early emergence of a leader-follower relationship that sustained into the remaining training sessions.

**Figure 3.**
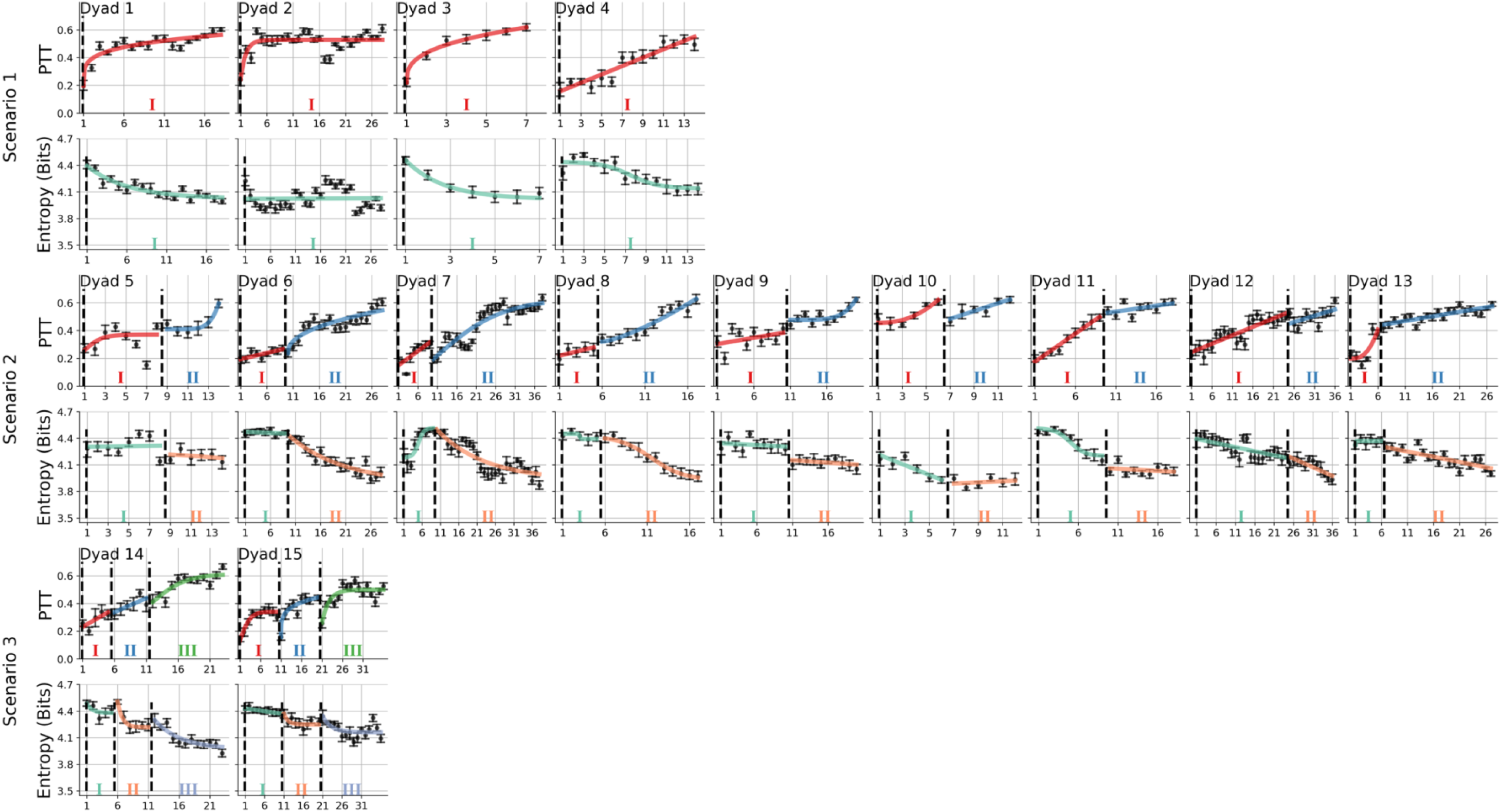
Mean proportion of time on task (PTT) and entropy across all training sessions for each dyad based on different scenarios. Different learning models are fitted to each training segment, which was determined based on the leader-follower relationship shown in Figure 2d. The predictions based on the model with the lowest Bayesian Information Criterion (BIC) are visualized as curves with different colors.

**Table 1.** Fitted model diagnostics for each segment.

| Dyad | Measure | I |  |  |  | II |  |  |  | III |  |  |  |
| --- | --- | --- | --- | --- | --- | --- | --- | --- | --- | --- | --- | --- | --- |
| | | Model | $R^2$ | $p$ | Sig. | Model | $R^2$ | $p$ | Sig. | Model | $R^2$ | $p$ | Sig. |
| 1 | PTT | Pow. | 0.67 | 0.003 | ** |  |  |  |  |  |  |  |  |
|  | Entropy | Exp. | 0.84 | 0.024 | * |  |  |  |  |  |  |  |  |
| 2 | PTT | Exp. | 0.33 | 0.007 | ** |  |  |  |  |  |  |  |  |
|  | Entropy | Lin. | < 0.01 | > 0.99 | n.s. |  |  |  |  |  |  |  |  |
| 3 | PTT | Pow. | 0.79 | 0.004 | ** |  |  |  |  |  |  |  |  |
|  | Entropy | Exp. | 0.99 | 0.002 | ** |  |  |  |  |  |  |  |  |
| 4 | PTT | Lin. | 0.61 | 0.039 | * |  |  |  |  |  |  |  |  |
|  | Entropy | Log. | 0.85 | 0.032 | * |  |  |  |  |  |  |  |  |
| 5 | PTT | Log. | 0.14 | 0.090 | n.s. | Pow. | 0.38 | 0.028 | * |  |  |  |  |
|  | Entropy | Lin. | < 0.01 | > 0.99 | n.s. | Lin. | 0.07 | 0.862 | n.s. |  |  |  |  |
| 6 | PTT | Lin. | 0.16 | 0.139 | n.s. | Pow. | 0.49 | 0.006 | ** |  |  |  |  |
|  | Entropy | Lin. | 0.07 | 0.786 | n.s. | Exp. | 0.87 | 0.018 | * |  |  |  |  |
| 7 | PTT | Lin. | 0.42 | 0.072 | n.s. | Log. | 0.71 | 0.001 | ** |  |  |  |  |
|  | Entropy | Log. | 0.84 | 0.061 | n.s. | Exp. | 0.79 | 0.021 | * |  |  |  |  |
| 8 | PTT | Lin. | 0.06 | 0.326 | n.s. | Pow. | 0.62 | 0.005 | ** |  |  |  |  |
|  | Entropy | Log. | 0.66 | 0.338 | n.s. | Log. | 0.98 | 0.005 | ** |  |  |  |  |
| 9 | PTT | Lin. | 0.07 | 0.213 | n.s. | Pow. | 0.39 | 0.016 | * |  |  |  |  |
|  | Entropy | Lin. | 0.04 | 0.872 | n.s. | Lin. | 0.18 | 0.527 | n.s. |  |  |  |  |
| 10 | PTT | Pow. | 0.42 | 0.024 | * | Lin. | 0.29 | 0.117 | n.s. |  |  |  |  |
|  | Entropy | Lin. | 0.79 | 0.190 | n.s. | Lin. | 0.07 | 0.849 | n.s. |  |  |  |  |
| 11 | PTT | Lin. | 0.69 | 0.041 | * | Lin. | 0.11 | 0.172 | n.s. |  |  |  |  |
|  | Entropy | Log. | 0.92 | 0.028 | * | Lin. | 0.06 | 0.817 | n.s. |  |  |  |  |
| 12 | PTT | Lin. | 0.45 | 0.041 | * | Lin. | 0.13 | 0.136 | n.s. |  |  |  |  |
|  | Entropy | Lin. | 0.53 | 0.157 | n.s. | Lin. | 0.67 | 0.172 | n.s. |  |  |  |  |
| 13 | PTT | Pow. | 0.43 | 0.022 | * | Lin. | 0.36 | 0.054 | n.s. |  |  |  |  |
|  | Entropy | Exp. | 0.54 | 0.382 | n.s. | Lin. | 0.60 | 0.146 | n.s. |  |  |  |  |
| 14 | PTT | Lin. | 0.12 | 0.215 | n.s. | Lin. | 0.12 | 0.199 | n.s. | Log. | 0.44 | 0.011 | * |
|  | Entropy | Exp. | 0.45 | 0.550 | n.s. | Exp. | 0.94 | 0.044 | * | Exp. | 0.83 | 0.044 | * |
| 15 | PTT | Log. | 0.53 | 0.009 | ** | Pow. | 0.59 | 0.008 | ** | Exp. | 0.45 | 0.008 | ** |
|  | Entropy | Lin. | 0.63 | 0.208 | n.s. | Exp. | 0.50 | 0.272 | n.s. | Exp. | 0.40 | 0.206 | n.s. |

For dyads in Scenario 2 (Single Role Switch; Dyads 5-13), the evaluation of model fitting diagnostics revealed two subcategories in learning patterns. First, for Dyads 5 to 9, the first training segment did not yield any statistically significant fit for any of the learning models, suggesting little to no systematic improvement in performance. After the leadership switch, the fitted learning models became significant with relatively high *R*^2^. This suggests that the new leader-follower configuration was more effective, allowing the dyad to develop a more stable and efficient coordination pattern. Curiously, most dyads within this group (except for Dyad 5) started the first segment with a linear trend, which was replaced by power law or logistic growth in the second segment. The contrasting learning patterns indicate that the dyads’ initial attempts at coordination were ineffective, resulting in minimal performance changes. Once the leadership role switched, learning trajectories followed non-linear trends (power law or logistic growth) with notable differences across dyads. Two dyads showed rapid initial improvement followed by a gradual slowdown (Dyads 6 and 7), while the other three dyads exhibited a slower initial learning phase that accelerated later (Dyads 5 and 8). This variation suggests that while some dyads quickly adapted to the new leader-follower dynamic before reaching a performance plateau, others required a longer adjustment period before experiencing a surge in improvement. In both cases, the leadership transition facilitated learning,

For the second subcategory, Dyads 10 to 13 showed significant learning in the first segment, indicating that the initial leader-follower relationship was effective in improving performance. However, after the role switch, none of the learning models significantly fit the data, suggesting that performance either stagnated or became more variable. Moreover, the models that best fit the first segment included power law (slow initial improvement followed by a rapid increase) and linear (consistent improvement). In contrast, only linear models were selected for the second segment, indicating that after the role switch, any observed changes in performance were gradual and lacked the accelerated learning observed earlier. This shift suggests that the second leader-follower configuration was less effective in driving further improvements.

Finally, Scenario 3 produced three distinct segments, reflecting a double leadership switch in which the original leader regained control in the final segment, leading to improved performance that reached the threshold to complete training. For Dyad 14, model fittings for the first two segments were non-significant, both following a linear pattern, suggesting little to no systematic improvement. This indicates that neither of the initial leader-follower configurations was effective in facilitating learning. However, in the final segment, when the leader-follower relationship reverted to its original configuration, learning became significant, following a logistic growth pattern characterized by rapid early improvement that slowed over time. This suggests that, although the initial leader-follower configuration did not facilitate effective learning, leadership switches and additional practice allowed the dyad to refine their coordination strategy. Over time, the dyad adapted to a more effective dynamic, ultimately leading to significant performance improvements in the final segment. For Dyad 15, all three segments yielded significant fits, with initial rapid performance improvement followed by a gradual slowdown. Despite the rapid growth, all segments started at around the same performance level (approximately 20% PTT) and finished at slightly higher performance than the previous segment. This suggests that while the dyad experienced consistent improvements across each segment, the gains were relatively small between transitions. This consistent pattern across segments could reflect an optimal coordination strategy being reached early on, with later improvements becoming more incremental.

### Leadership Dynamics and Coordination Stability

The analysis of learning trajectories revealed distinct patterns in how leader-follower dynamics influenced learning over time. To further explore these dynamics, Shannon entropy^30^ was used to quantify the coordination stability to reveal the factors driving the leadership switch. Shannon entropy is a measure of uncertainty or unpredictability in a dyad system, quantifying the level of disorder or variability in the coordination over time. Entropy was calculated at both the system level (using the relative phase between individuals) and the individual level (based on each participant’s phase), providing a means to assess coordination stability. A decrease in system entropy may reflect the emergence of a stable coordination strategy, while changes in individual entropy could indicate adaptive adjustments in response to shifting leader-follower roles.

#### System Stability

On a system level, entropy trends across sessions closely mirrored changes in performance resulting from learning (Figure 3b, Table 1). For dyads in Scenario 1 (Persistent Leadership), system entropy decreased over training sessions, indicating increasing coordination stability. This reduction in entropy reflects the dyadic system’s progression toward more predictable and organized movement patterns, which coincided with improved performance over time. For dyads in Scenario 2 (Single Role Switch), those without a statistically significant model fit for the first segment (Dyads 5 to 9) also showed non-significant changes in entropy. This suggests that these dyads struggled to initially establish an effective coordination strategy, resulting in persistent system instability and limited performance gains. In the second segment, however, entropy trends were generally significant and aligned with improved performance, indicating that once a successful coordination strategy emerged, system stability increased alongside performance improvements. In contrast, for those with a statistically significant model fit for the first segment (Dyads 10 to 13), only one out of the four dyads yielded a significant entropy fit for the first segment. This discrepancy suggests that improved performance may not always coincide with greater stability in coordination. This could be attributed to partial stability in coordination, where performance gains are achieved through strategies that remain variable or inconsistent. Finally, the aforementioned observations for Scenario 2 can also be applied to Scenario 3 where the system entropy either reflects the attainment of a stable coordination strategy (Dyad 14) or a lack thereof (Dyad 15).

#### Individual Stability

To examine how individual stability shapes leadership dynamics, individual entropy was calculated for each partner (referenced as top and bottom dots) in each trial. Across all trials within a session, various entropy-related features were extracted, such as the mean, standard deviation, minimum, and maximum entropy for each dot. Because the proportion of trials in which the top and bottom dots were leaders are mathematically complementary within each session (e.g., if the top dot was the leader in 60% of trials, the bottom dot was the leader in the remaining 40%), we chose the top dot’s leadership percentage as the outcome variable. Entropy features were then used as predictors for the proportion of trials in which the top dot was the leader, modeled using a series of generalized linear mixed models (GLMMs) with a beta distribution and a logit link function. All possible combinations of the eight predictors were tested (2^8^ – 1 = 255) and compared using the Akaike information criterion (AIC)^31^ to identify the best-fitting model, which yielded:

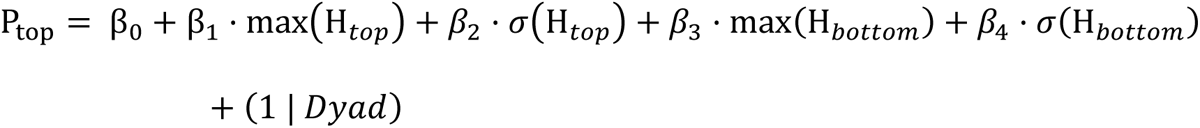

Where P_top_ is the proportion of trials in which the top dot assumes the leader role during a session. H_*top*_ and H_*bottom*_ are the entropy of the top and bottom dots, respectively. *σ* denotes the standard deviation of entropy, and max refers to the maximum entropy observed within a session.

The model provides a strong fit to the data (Efron *R*^2^ = 0.357). There was a significant negative effect of the standard deviation of the bottom dot’s entropy (*β*_4_ = 6.57, SE = 1.44, *p* < 0.001) on P_top_. While the standard deviation of the top dot’s entropy was included in the final model, its parameter was not statistically significant (*β*_2_= -2.34, SE = 1.58, *p* = 0.14). Removing this term led to a slightly higher AIC (ΔAIC = 0.21), suggesting that the term may offer additional explanatory value when included with the other predictors, despite its lack of statistical significance. The complementary effects of the top and bottom dots’ entropy variability suggest that the emergence of leadership roles depends on the asymmetry of movement variability between the individuals. Specifically, higher variability in the follower (bottom dot) may prompt the system to stabilize around the more consistent individual (top dot), reinforcing its leadership. In contrast, excessive variability in the leader (top dot) may undermine its stability and reduce the likelihood of being perceived as the reference point for coordination. This balance of variability across agents underscores the importance of differentiated roles in achieving stable coordination.

A similar complementary asymmetry between the individuals in a dyad also manifested in maximum entropy. There was a significant positive effect of the top dot’s maximum entropy (*β*_1_ = 3.87, SE = 0.49, *p* < 0.001) and a significant negative effect of the bottom dot’s maximum entropy (*β*_3_ = -1.09, SE = 0.41, *p* < 0.01). In the present context, maximum entropy reflects momentary bursts of instability in an otherwise more stable signal, as indicated by the patterns observed in the entropy’s standard deviations. For the top dot, such bursts may function as assertive or corrective movements that reorient the system around them, facilitating leadership through a more dominant movement pattern. Conversely, bursts of instability in the bottom dot may themselves act as mechanisms for leadership emergence, prompting the system to reorganize around the bottom dot instead. This implies that momentary instability can support leadership for either dot, depending on the broader context: for the top dot, it enhances dominance; for the bottom dot, it may signal a shift in control. These findings underscore the importance of temporal instability as a flexible and context-dependent mechanism in leader-follower dynamics.

## Discussion

Research on human and animal behaviors has traditionally attributed leadership in coordinated actions to intrinsic individual characteristics^6,9^, pre-defined social structures^7,8^, or cognitive functions^10^. Findings from the current study demonstrated that leadership is not merely an inherent trait or predetermined role, but rather an emergent property of the dyad system. It is dynamically modulated by the interplay between individual actors and the system’s stability, which, in turn, influences overall system performance and improvement. This dynamic perspective suggests that leadership is not solely dictated by fixed attributes but fluctuates based on contextual demands and the ongoing interactions between individuals in the system. As the system evolves, individual adaptations and mutual adjustments shape the leader-follower relationship, reinforcing or shifting leadership roles as needed. Importantly, our findings indicate that effective coordination does not necessarily require a consistently dominant leader; instead, transient leadership shifts may facilitate adaptability, particularly in tasks requiring high levels of collaboration and synchrony.

A critical factor in the learning process was the ability of one individual to establish a stable reference movement. Dyads with an effective leader emerging early exhibited smoother learning trajectories, suggesting that an early established leader-follower dynamic facilitates learning. Conversely, when the initial leader struggled to maintain a consistent rhythm or the follower failed to adjust, learning was either delayed or plateaued. In such cases, leadership transitions sometimes unlocked progress but, in other instances, disrupted already-established coordination patterns, leading to temporary setbacks. These findings align with previous research that suggests that stable leadership in joint action tasks is key to optimizing dyadic performance^32^. High partner variability can hinder individual performance, yet predictable movements, particularly those emerging from stable leader-follower dynamics, can offset these negative effects. This predictability enhances flexibility and resilience within the dyad, improving overall coordination^33^. This finding underscores the importance of synchronized behaviors in joint tasks, where participants adapt their actions based on their partner’s movements, demonstrating an ability to utilize their own actions to respond to others’ actions^34^.

Coordination patterns are prevalent across natural systems, influencing both individual and group behaviors. The foundational theory of coordination dynamics^35,36^ posits that such patterns emerge through self-organization rather than fixed programming, driven by interactions among system components, whether neurons, limbs, or individuals. Perceptual information plays a central role in shaping these dynamics, acting as the medium through which components interact and adjust^22,37,38^. In the present study, dyads relied exclusively on shared visual information to learn the 90° coordination. The oscillating dots provided continuous, real-time information about a partner’s direction and position, enabling the detection of relative phase^26^ and allowing for continuous mutual adjustments. This form of bidirectional visual coupling invited both intra- and inter-personal variability into the coordination process, a dynamic supported by our entropy analyses.

While bidirectional visual coupling might imply symmetrical variability between partners, our results revealed a pronounced asymmetry in movement variability that reflects the functional distinction between leader and follower roles. Such asymmetry was likely shaped by the presence of the auditory metronome, which constrained one partner’s behavior more than the other’s. Acting as a fixed-frequency attractor, the metronome helped stabilize the system but also introduced a coordination challenge: achieving a 90° offset required each individual to continuously align their movement not only with their partner but also with the external beat. Our data suggest that dyads spontaneously adopted a division of labor strategy, where one individual (the leader) prioritized synchronization with the metronome, while the other (the follower) adjusted based on the partner’s visual cues. This strategy not only facilitated dyadic performance but also accounted for the observed differences in stability. The leader’s proximity to the metronome as an attractor promoted consistency, whereas the follower’s reliance on adapting to an inherently variable partner induced greater motor variability. Interestingly, leaders occasionally exhibited brief bursts of instability, which may reflect lapses in attentional focus due to the compelling nature of visual information^39^. In moments where attention shifted from the auditory to the visual channel, alignment with the metronome temporarily degraded, followed by recalibration. These observations underscore the multi-modal nature of coordination dynamics: While auditory information stabilized performance, visual information enabled dynamic correction and adaptation, potentially at the cost of stability. More broadly, this finding highlights the malleability of coordination dynamics in response to perceptual context. Modifying the nature or availability of perceptual information, for instance, introducing a haptic connection between partners, could fundamentally restructure leader-follower dynamics and the distribution of variability^40^. Such shifts could open new pathways for studying and designing collaborative systems, especially those that rely on shared sensorimotor contingencies.

Anchored by stable reference signals and adaptive responses, this coordination strategy mirrors principles in complex adaptive systems, where roles such as leader and follower emerge not through explicit designation, but through interactions between component states and environmental demands. For instance, in a study of pairs of foraging animals^6^, individuals with different energetic states or levels of predation risk spontaneously adopted distinct roles, where the hungrier individual emerged as the pace-maker, driving decisions about when and where to forage, while the other adjusted accordingly. Drawing a parallel to our findings, the brief kinematic bursts observed in the leader’s movement may represent behavioral expressions of such pace- making. These transient deviations from a stable rhythm could be interpreted as initiation signals, that is, moments where the leader asserts directional control to reorganize or reinforce the dyad’s coordination strategy. Just as the foraging leader uses energetic urgency to prompt movement, our task leader may use temporal or spatial drift as a mechanism to re-anchor the dyad’s phase relationship. Similarly, informational asymmetries, such as one partner having clearer access to or confidence in the metronome, could drive these bursts, turning abstract role differentiation into a measurable motor signal. From this perspective, kinematic bursts are not simply noise or error, but rather an adaptive feature of dyadic coordination. They embody the leader’s moment-to-moment recalibration function within the system, allowing the dyad to remain flexible in the face of uncertainty. This behavioral interpretation may offer a concrete bridge between ecological models of group coordination and embodied movement studies in humans, providing a new lens on how leadership can emerge and fluctuate in real-time interactions.

In conclusion, this study highlights how coordination and learning in joint action are not simply products of individual traits or imposed hierarchies, but rather the emergent result of dynamic interactions between individuals shaped by informational constraints and performance stability. Through the lens of complementary asymmetric stability, we revealed that momentary kinematic bursts in leaders and sustained variability in followers form a functional pattern that supports adaptive coordination. These findings challenge traditional, fixed-role conceptions of leadership by demonstrating how leadership can arise fluidly and temporarily in response to the system’s needs. Moreover, by grounding our interpretation in coordination dynamics and perceptual coupling, we offer a mechanistic account of how sensorimotor information shapes emergent roles in real-time interactions. Taken together, this work contributes to a growing understanding of leadership as a distributed, relational property, and offers a framework for studying how human systems can flexibly self-organize to meet shared goals under varying constraints.

## Methods

### Participants

Thirty participants (18 – 35 years old) were recruited from the University of Wyoming community and then randomly paired to form 15 dyads. All participants were right-handed and had normal or corrected-to-normal vision. This study was approved by the University of Wyoming Institutional Review Board (IRB). Informed consent was obtained from all participants before participation following the Declaration of Helsinki.

### Stimuli and Apparatus

Figure 1a illustrates the experimental setup. Participants sat at a desk arranged parallel to and adjacent to a column in the testing room. The column, along with an adjoining wall, separated the two seats, blocking participants’ views of each other. A shared 17.3-inch laptop display was positioned at eye level between the participants. Two Logitech Extreme 3D Pro joysticks were connected to the laptop, and each participant used their dominant hand to control a joystick, moving one of two white dots displayed on the screen to produce a 90° coordination pattern (Figure 1b). The white dots, each 1 cm in diameter, moved horizontally along parallel paths on the screen, with one dot positioned above the other. The horizontal path length was 25 cm, and the vertical distance between the paths was 7.5 cm. Dot positions were recorded at a sampling frequency of 120 Hz. Participants could only view the movements of the two dots on the shared display, as their partner’s limb movements were occluded by the wall. The distance from the participant’s eyes to the center of the screen was approximately 75 cm. The participants were instructed to not talk to each other during the sessions.

### Procedures

Each dyad completed 1 to 2 training sessions per day until their average performance of the 90° relative phase in a single session reached 60% Proportion of Time on Task (PTT), the primary measure of coordination performance in this study. Each training session comprised 20 trials, with each trial lasting 20 seconds. At the beginning of each session and after every five trials, participants were shown an 8-second visual demonstration of the 90° coordination pattern at 0.75 Hz as a reminder. During training, a metronome set to 0.75 Hz provided auditory guidance for frequency-specific learning of the 90° coordination pattern for the first 10 seconds of each trial. Visual feedback was also incorporated, where the dots turned green when the relative phase fell within the target range of 90° ± 20°, reverting to white when the phase deviated outside this range.

### Data Analysis

Kinematic data were processed using TAT-HUM, a human kinematic analysis toolkit in Python^29^. Raw position data, *x*, were first filtered with a low-pass Butterworth filter with a 120 Hz sampling frequency and a 10 Hz cutoff frequency. Velocity, *v*, was derived using a finite differences method. The phase at each time point was determined as the angle of the velocity-position vector in the phase plane:

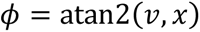

Using atan2 ensures that the phase is correctly assigned to the appropriate quadrant of the circular phase plane, yielding values in the range of [−*π*, *π*]. To address the discontinuity at around 2*π*, phase unwrapping was performed. When transitions occurred where the previous phase was near 2*π* and the current phase was near 0, 2*π* was added to the current phase to ensure continuity. This process resulted in a continuous phase representation for each trial (Figure 2a).

The phase represents the dot’s position within its oscillation cycle and can be used to determine the relationship between the two dots. For example, if the phase magnitude of the top dot is consistently greater than that of the bottom dot during a trial, it indicates that the top dot was leading while the bottom dot followed in producing the coordination pattern. To quantify this relationship, the *phase difference* (Δ*ϕ*) between the top and bottom dots was computed at each timestamp, *t* (Figure 2b):

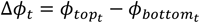

Task performance was evaluated using PTT, which requires folding the phase difference at 180° to ensure that the leadership (top vs. bottom) did not affect the measure (Figure 2c). PTT was defined as the proportion of time during which the relative phase remained within a ±20° error band around the target phase.

To identify the leader-follower relationship within a dyad, phase difference was used. For each trial, a one sample t-test was performed to compare the phase difference with 0 at each timestamp. If the relative phase was significantly greater than 0, the top dot was identified as the leader, and vice versa for the bottom dot if the phase difference was significantly less than 0. If the t-test did not yield a significant result, neither dot was considered the leader, and the trial was removed from subsequent analysis. Across all training sessions and all dyads, there were only 18 trials (or 0.25% of total trials) in which neither individual in the dyad was the leader. Subsequently, the proportion of trials in which one individual was the leader was calculated for each session, and training sessions were then segmented based on whether one individual consistently led for three or more consecutive sessions (Figure 2d).

Different learning models were fitted to the PTT data for each training segment to explore how the leader-follower relationship affects motor learning. Four learning models were used, including linear, power law, logistic growth, and exponential decay (Table 2). For each training segment, each model was fitted to the PTT of every trial and session. Bayesian information criterion (BIC) was used for model selection, where the model that yields the smallest BIC score was selected and the model’s statistical significance (*p*-value) and goodness-of-fit (*R*^2^) were reported. BIC is used because some fitted segments may contain fewer data points and BIC’s stronger penalty for model complexity makes it appropriate to avoid overfitting.

**Table 2.** Fitted Learning Models for the Training PTT Data.

| **Model** | **Expression** | **Free Parameters** |
| --- | --- | --- |
| Linear | $PTT = a \cdot t + b$ | $a, b$ |
| Power Law | $PTT = a \cdot t^b + c$ | $a, b, c$ |
| Logistic Growth | $PTT = P_0 + \frac{P_{\max} - P_0}{1 + e^{-kt}}$ | $k, P_0, P_{\max}$ |
| Exponential Decay | $PTT = P_0 + (P_{\max} - P_0)(1 - e^{-kt})$ | $k, P_0, P_{\max}$ |

To further evaluate the relationship between leadership dynamics and training performance, the Shannon entropy was derived for the individual phases of the top and bottom dots as well as their relative phase. The individual phases and relative phase were wrapped around π to derive their respective frequency distributions based on 100 bins, which were used to compute their respective entropy:

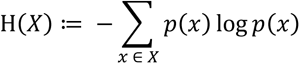

Entropy derived from the relative phase captures the dyadic system’s coordination stability, reflecting the degree of predictability or variability in the movement patterns between the two individuals. Similar to PTT, the system entropy was segmented based on the leadership dynamics where each segment was fitted to one of the four learning models (Table 2) and the best model was selected based on BIC.

Entropy derived from individual phases describes the degree of unpredictability or variability in each individual’s movement, providing insight into the stability and adaptability of their behavior. High entropy in an individual’s movement may indicate periods of instability or rapid changes, while low entropy may suggest more consistent and controlled movement patterns. To examine the relationship between individual entropy and leadership dynamics, different features of the individual entropy were computed, including its mean, standard deviation, minimum, and maximum, for each session. These features were used as predictors in a series of GLMMs with a beta distribution and a logit link function to predict the proportion of trials where the top dot was the leader. All possible combinations of features were tested to identify the best-fitting model for the data. Model selection was performed using AIC, which is more suitable for complex models with hierarchical structures. This approach allows for a more accurate assessment of how variations in individual entropy relate to the likelihood of leadership emergence in the dyadic system.

## Data Availability

The data supporting the findings of this study are available at https://osf.io/upxcz/.

## Code Availability

The analysis code supporting the findings of this study is available on GitHub at https://github.com/xywang01/dyad-leadership.

## Acknowledgment

The authors would like to thank Shaiyan Keshvari for his helpful comments and discussion.

